# Rapid adaptation with gene flow via a reservoir of chromosomal inversion variation?

**DOI:** 10.1101/150771

**Authors:** Qixin He, L. Lacey Knowles

## Abstract

The increased recognition of frequent divergence with gene flow has renewed interest in chromosomal inversions as a source for promoting adaptive divergence. Inversions can suppress recombination between heterokaryotypes so that local adapted inversions will be protected from introgression with the migrants. However, we do not have a clear understanding of the conditions for which adaptive divergence is more or less likely to be promoted by inversions when the availability of inversion variation is considered. Standing genetic variation, as opposed to new mutations, could offer a quick way to respond to sudden environmental changes, making it a likely avenue for rapid adaptation. For a scenario of secondary contact between locally-adapted populations, we might intuit that standing inversion variation would predominate over new inversion mutations in maintaining local divergence. Our results show that this is not always the case. Maladaptive gene flow, as both a demographic parameter and the cause for selection that favors locally-adapted inversions, differentiates the dynamics of standing inversion variation from that of segregating point mutations. Counterintuitively, in general, standing inversion variation will be less important to the adaptation than new inversions under the demographic and genetic conditions that are more conducive to adaptive divergence via inversions.

## Introduction

Allopatric populations usually accumulate locally adaptive alleles after a period of environmental change (Nosil et al. 2009a; Papadopulos et al. 2011). How fast can the adaptive divergence occur? The two most important factors, sources of variation and the probability of fixation of favorable mutations, have been thoroughly discussed from classical population genetic theories (e.g., Ewens 2004; Fisher 1930; Kimura 1983; Orr 1998) to empirical data on patterns of genetic variation (e.g., Bradshaw et al. 1995; Colosimo et al. 2005; Karasov et al. 2010). Less is clear about the mechanisms that maintain these adaptive loci in the face of maladaptive gene flow after secondary contact, which is common at the early stage of adaptive divergence (Nosil et al. 2009b). Specifically, gene flow from ecologically dissimilar populations will dilute locally-adapted loci and disrupt the combinations of alleles by recombination. Under such scenario, any mechanism that can lower the effective gene flow rate or protect the good combination of alleles from shuffling with bad alleles in recombination will be preferred in adaptation (Yeaman and Whitlock 2011). Rearrangements in chromosomes – inversions – can serve as such a mechanism because they can suppress recombination between heterokaryotypes so that local chromosomes carrying the adaptive alleles within an inversion will be protected from introgression of the maladapted genes carried by the migrants (Kirkpatrick 2011; Kirkpatrick and Barton 2006; Manoukis et al. 2008; Navarro and Barton 2003; Noor et al. 2001; Rieseberg 2001).

Empirical evidence has shown that many adaptive loci are associated with inversions, especially complex traits such as wing patterns (Joron et al. 2011), diapause timing (Feder et al. 2003a) and annual/perennial life-history shift (Lowry and Willis 2010). However, it is not clear whether these inversions become established in the population because of the maladaptive gene flow. Although theoretically possible, and despite the appeal of such a hypothesis, we don’t have a clear understanding of the conditions (genetic or demographic) for which adaptive divergence is more or less likely to be promoted by inversions. Similar to adaptive point mutations, the rate of adaptation via locally-adapted inversions is determined by the availability of inversion variation (i.e., either new inversion mutations or pre-exisiting standing variation of inversions) and the probability of establishment of favorable inversions. Both aspects were modeled or simulated separately in several studies (Feder et al. 2011; Kirkpatrick and Barton 2006; Manoukis et al. 2008). What is missing for evaluating the contribution of inversions to adaptation is critical information on the rate of adaptation that considers the probability of inversions capturing a locally-adapted genotype, as well as the likely contribution of standing inversion variation versus new inversion mutations.

Here we develop a theory for rapid local adaptation under gene flow via chromosomal inversions such that we are able to predict when (i.e., genetic or demographic conditions) the rate of adaptation by inversions will be higher. Moreover, we can evaluate the likelihood of standing inversion variation contributing to adaptive divergence. By expanding the repertoire of models of local adaptation, our work contributes to a growing body of work for predicting when different mechanisms are likely to promote rapid evolution (e.g., Hermisson and Pennings 2005; Kirkpatrick and Barton 2006; Przeworski et al. 2005; Scoville and Pfrender 2010). Moreover, by focusing on the potential contribution of new mutational input versus standing genetic variation, the general rules derived from the developed theory takes on special significance given the difficulty for such distinctions based on empirical evaluations of molecular data (reviewed in Barrett and Schluter 2008).

Similar to standing variation of point mutations that facilitate rapid adaptation under sudden environmental change (Orr and Betancourt 2001; Przeworski et al. 2005) and bottlenecks (Hermisson and Pennings 2005; Orr and Unckless 2008), standing inversion variation can be readily established in the population without a prolonged waiting time for the occurrence of an inversion and will suffer less random loss compared to new inversions if the mean frequency is larger than 1/2N. However, unlike point mutations, inversions do not confer a fitness difference directly. Instead, they reduce recombination cost and create linkage disequilibrium among selected loci. Therefore, the chance of local adaptation from a new inversion through indirect selection might be very low considering that the probability of an inversion mutation capturing coadapted genotypes would be smaller after the onset of gene flow, let alone the possible stochastic loss of the single mutation. This contrasts with standing inversion variation (i.e., inversions that captures good combinations of alleles before the onset of gene flow). Moreover, standing inversion variation will have less chance of harboring deleterious mutations because they have been under purifying selection before the onset of gene flow. These factors might enhance the importance of standing inversion variation in the scenario of secondary contact.

Our key finding is that when inversions facilitate divergence with gene flow, higher gene flow increases the contribution from standing inversion variation. Yet, under the demographic and genetic conditions that are more conducive to adaptive divergence via inversions, new inversions become a more important source. We discuss how this counterintuitive result (and one that differs from a recent study; Feder et al. 2011) can only be understood by explicitly considering the dynamics of adaptation from inversion variation, highlighting the importance and utility of analytical models for studying adaptation. By considering a broad range of selective values of alleles, instead of assuming weak selection (Kirkpatrick and Barton 2006), our model and simulation also include predictions about (i) the characteristics of inversions contributing to adaptation (e.g., the selective benefit of alleles and its relationship with migration and number of loci involved) and (ii) conditions when the relative importance of standing inversion variation as a source of maintaining adaptive divergence might be increased.

## Models and Methods

Consider a situation in which a peripheral population is receiving maladaptive gene flow from the central population across a heterogeneous environment such that alleles at two (or more) loci confer a selective benefit in the peripheral population but are maladaptive in the central population (Fig. 1). If the genetics of local adaptation is based on more than one locus, the maintenance of adapted alleles in the gene flow will depend not only on the selective benefit of an allele, but also on whether recombination will break up coadapted alleles (i.e., produce genotypes with a combination of adaptive and maladaptive alleles). If a chromosomal inversion captures the locally adapted alleles, with the introduction of maladapted alleles by gene flow (Fig. 1), there is a selective advantage to adapted alleles captured in an inversion because of suppressed recombination, whereas in the standard (i.e., non-inverted) chromosome locally adapted alleles can freely recombine with maladapted migrant alleles in heterokaryotypes, thereby breaking up locally adapted genotypes (Kirkpatrick and Barton 2006).

**Fig. 1.**
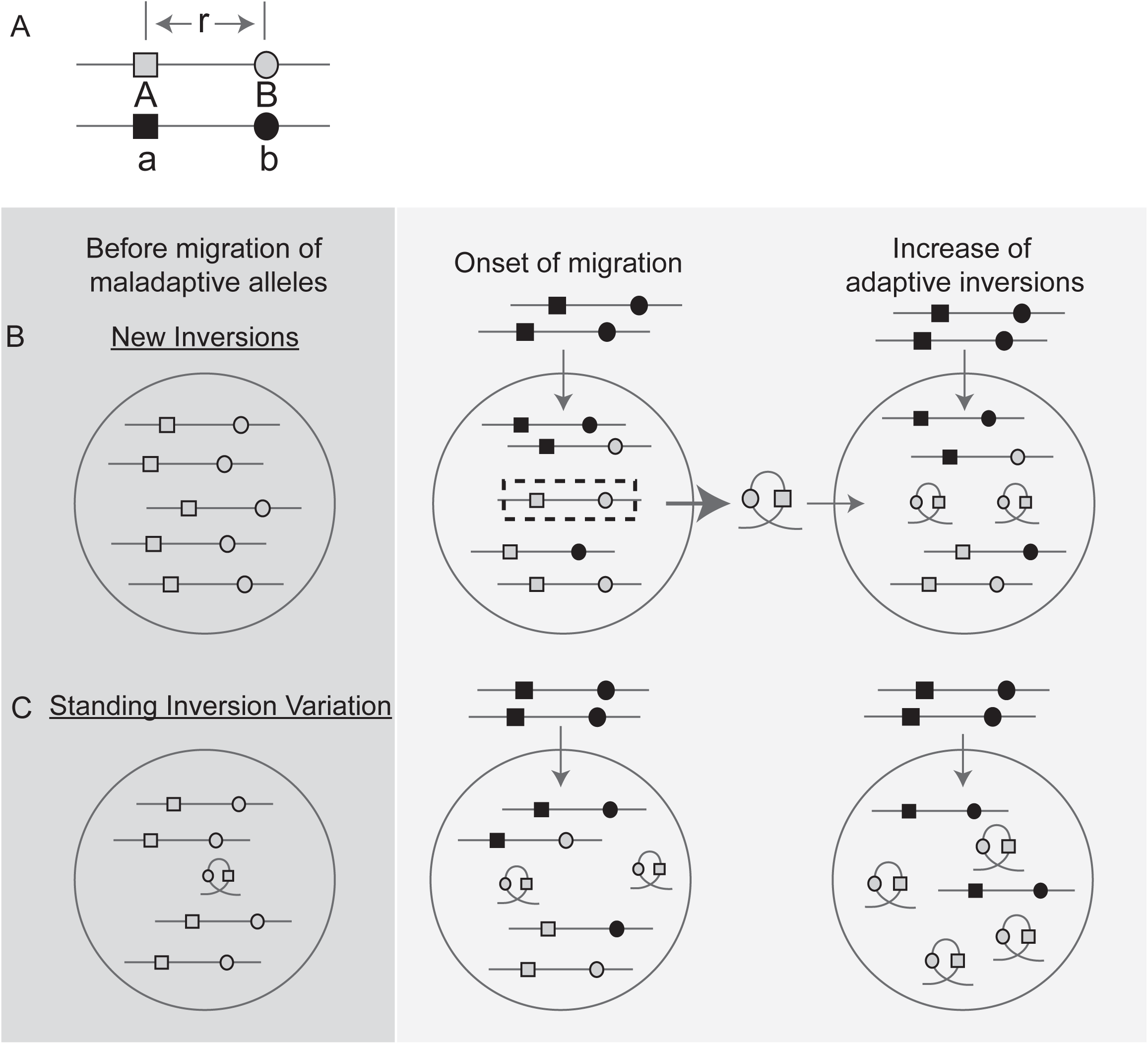
Illustration of the processes involved in adaptation from inversions under divergence with gene flow. (A) For two loci (shown by a square and a circle), alleles *A* and *B* (shown in grey) are locally adapted compared with the maladapted alleles, *a* and *b* (shown in black); the two loci are linked on the same chromosome with a recombination fraction, *r*. (B) Adaptation from new inversion and (C) standing inversion variation when divergence occurs with gene flow. See method section for the explanation of the process.

Based on the scenario described above, we focus on comparing the dynamics and probabilities of maintenance of divergence from new inversions and standing inversion variation. Consider the simplest scenario of a single inversion mutation that captures locally coadapted alleles of two loci in a diploid population, where alleles *A* and *B* each have a homozygous fitness advantage *s* with dominance coefficient *h* over the maladapted alleles *a* and *b* from a different population (Fig. 1A). The two loci are linked on the same chromosome with recombination fraction *r* (Fig. 1A). These loci have independent influence on the individual fitness (i.e., multiplicative fitness is assumed, meaning no epistasis, Table 1). Before onset of migration, the populations are monomorphic with haplotype *AB*. With migration, maladapted alleles (*ab*, shown in black) replace *m* proportion of the locally adapted *AB* individuals each generation, and forming recombinants (*Ab*, *aB*) with homokaryotes (i.e., with the standard, non-inverted chromosome). We define inversions mutations that capture the coadapted genotype occurs after gene flow started as new inversions (NI, Fig. 1B), and the ones that segregate in the population before gene flow as standing inversion variation (SIV, Fig. 1C). The standing inversion variations are selectively neutral until the start of gene flow because individual fitness is determined only by the alleles. Denoted as *AB*\*, recombination between inversions and other standard karyotypes is suppressed. Because migrants can only form recombinants with standard chromosomes, the inversion *AB*\* will become more advantageous (if it survives the initial stochastic loss) as the proportion of recombinants, *Ab* and *aB,* is built up. When adaptation from inversions is successful, all the other genotypes, *AB*, *Ab*, *aB*, will be replaced by *AB*\* with *ab* left in the population if gene flow continues (in this study this stage is referred to as establishment of the inversion).

**Table 1.**
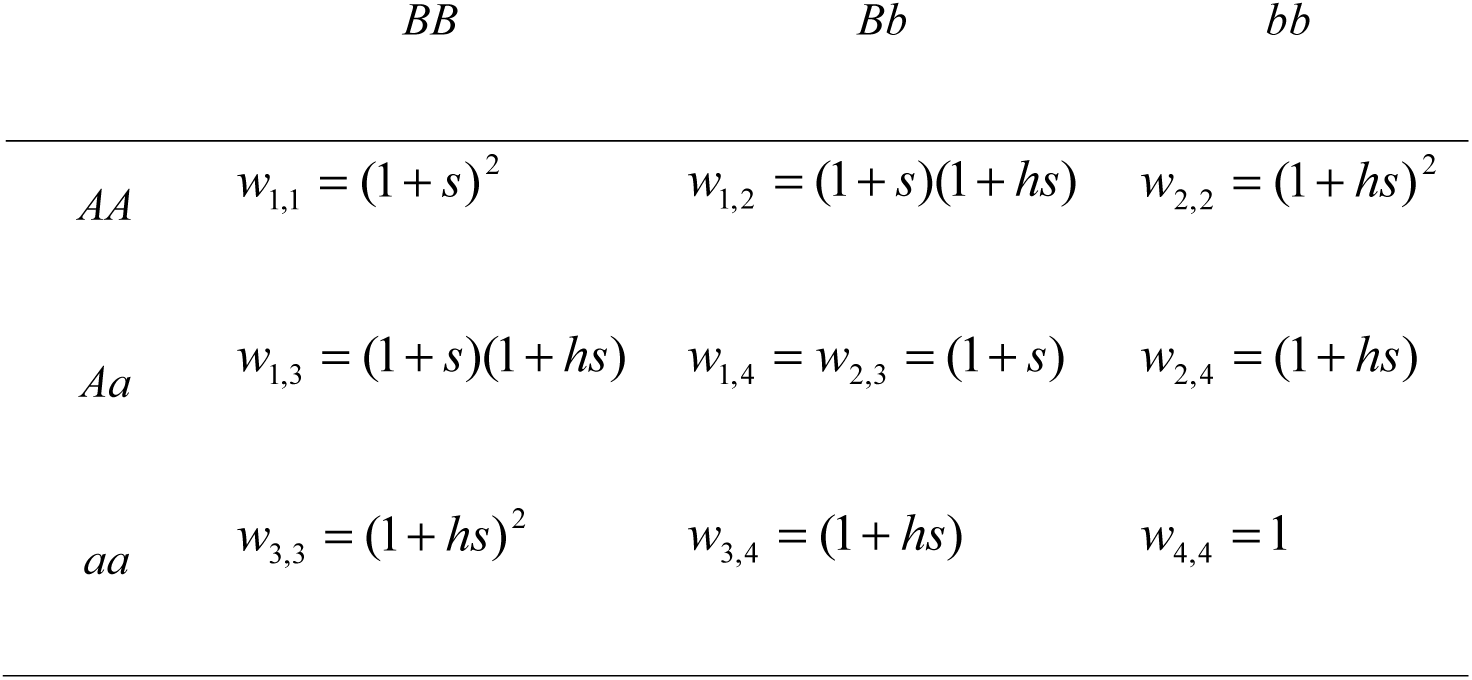
Fitness of offspring from different parental genotypes, where the haplotype *AB* is locally adapted (see Fig. 1) assuming loci have independent fitness effects (i.e., multiplicative fitness, or no epistasis).

The rate of spreading of an inversion depends upon the following parameters in a deterministic model: gene flow rates, allele effect sizes, number of locally-adapted loci and rate of recombination between them. When the allele effect size is small (i.e., s<<1), it can be omitted from the analytical approximation (see Eq 1. in Kirkpatrick and Barton 2006). However, if allele effect size is not negligible, linkage disequilibrium (LD) between adapted loci needs to be considered to calculate the frequency of genotypes. Here we relaxed the assumption, and derived analytical approximations to examine the relationship between gene flow (*m*) and allele effect size (*s*) in determining the rate of adaptation via inversions. Comparisons between the probabilities of adaptation from new inversions with that from standing inversion variation are also evaluated in the context of the availability of these two sources under different parameter spaces.

All analytical equations were tested using time-forward simulations (Mathematica and Matlab code available at Dryad, to add upon acceptance). At each generation, with a population of *N* diploid hermaphrodically reproducing individuals (*N_e_* = 5000), migration (*m*) occurs first, followed by selection of individuals according to their fitnesses (Table 1); recombination occurs at rate *r* as gametes are formed meiotically, and then the next generation of dipoloids is randomly drawn from the pool of gametes. Each individual in a population is represented as two linear chromosomes of *n* loci with same allele effect (*s*) and no dominance (*h* =0.5). Recombination is suppressed in heterokaryotypes. Free recombination is assumed between loci (i.e., *r* = 0.5), except for simulations that explore the effects of specific parameter values. In new inversion case, migration is allowed to occur until the population reaches migration-selection-drift balance. New chromosomal inversions are introduced at a mutation rate, *μ,* of 10^-7^ per gamete per generation in a population at migration-selection-drift balance. The inversion is tracked and generation time is recorded until when the inversion either goes extinct or replaces all the other adapted genotypes (i.e., selected loci in non-inverted chromosomes). In the standing inversion variation case, the starting frequency of the standing inversion variation for each simulation is selected at random from the expected distribution of neutral segregating inversions under the same demographic settings (generated by forward simulation with over 10,000 generations). The inversions are again tracked until either their loss or establishment. To determine whether new inversions or standing inversion variation is more likely to contribute to rapid adaptation, a population allowed for both new inversions and standing inversion variation is generated (as described above). The proportion of simulations in which each of the two sources of adaptive variation are either lost or established is quantified. 10,000 replicates were run for each set of parameter values.

## Results

### Selective advantage of a new inversion

In the simplest scenario of a single inversion mutation that captures locally coadaptive alleles of two loci in a diploid population (Fig. 1), gametes *AB*, *Ab*, *aB* and *ab* have the genotype frequencies *x*_1_,*x*_2_,*x*_3_,*x*_4_. The change of gametic frequencies will be 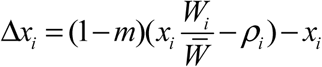 (Li and Nei 1974), where *ρ_i_* is the change in frequency of *x_i_* attributable to recombination with either locally coadapted or maladaptive alleles. It is defined as 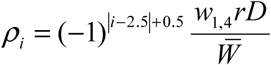 (Lewontin and Kojima 1960), where *w*_1,4_ is the fitness of double heterozygotes *A*/*a B*/*b* (Table 1) and *D* is the coefficient of LD, *x*_1_*x*_4_ – *x*_2_*x*_3_. At migration-selection balance, where 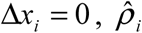 can be expressed as 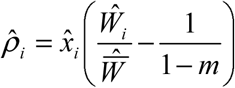.

When an inversion captures coadapted alleles, denoted as *AB*\* (Fig. 1), its frequency will increase in the next generation as a function of 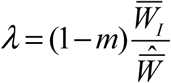 (Kirkpatrick and Barton 2006), where 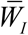 is the average fitness of individuals with an inverted chromosome and 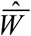 is the average fitness of the population at equilibrium, which is determined by the frequencies of the four gametes and the fitness of each genotype (Table 1). The initial increase in the frequency of a new inversion, *λ*, is therefore proportional to the decrease of the frequency of the coadapted genotype attributed to recombination scaled by gene flow,

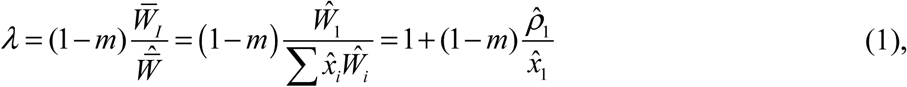

where 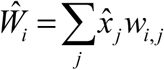. The inversion will always be favored by selection (i.e., 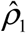 will be positive) as long as *s* is large enough such that locally adapted alleles *A* and *B* can withstand the swamping effect of migration of maladapted alleles *a* and *b* (i.e., *s* > *m*/(1–*m*) for *s*<<1; there is no simple approximation if *s* is larger). Adaptation occurs by the increase in the frequency of the inversion, which will reduce the effective gene flow rate and elevate the mean fitness of the population.

### Probability of establishment of a new adaptive inversion

The probability of establishment of a new adaptive inversion (*f_NI_*) is determined by its selective advantage in the first few generations. Using a branching process approach, classical work showed that it is approximately twice its initial selective advantage weighted by the reproductive variance (i.e., 2(*λ*-1)/*λ*, Haldane 1927; Kimura 1957). Hence, a new adaptive inversion will be established in the population with the probability

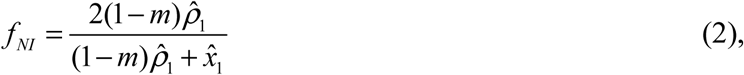

assuming the number of offspring per parent is Poisson distributed (such that the reproductive variance of inversions equals *λ*) under a Wright-Fisher model. Using numerical approximations to explore the establishment probability of *AB*\* under different combinations of *m* and *s* (Fig. 2A), we show that it is not just the rate of migration (Kirkpatrick and Barton 2006), but that the allele effect size is also important. The greatest probability of establishment of a new inversion, *f_NI_*, occurs when allele effect size is the smallest. This can be intuitively understood by considering that when locally coadapted alleles are captured by an inversion, they never suffer the disadvantage of being found with maladapted immigrant alleles because of suppressed recombination. However, the selective advantage of inversions decreases as the allele effect sizes increase because 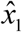, the frequency of the coadapted genotype *AB* on the standard (i.e., non-inverted) chromosome also increases when the allele effect size increases (Eq. 1). Overall, the migration rate, *m*, is the primary determinant of the probability of establishment of a new inversion (Fig. 2A), having a larger effect on the probability of establishment of the new inversion than the allele effect size. Nevertheless, there is a limit to which migration can facilitate the adaptation from inversions and this limit is determined by the effect size associated with the contained alleles. Specifically, we show that the equilibrium frequency of inversions, 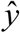, decreases at high gene flow rates,

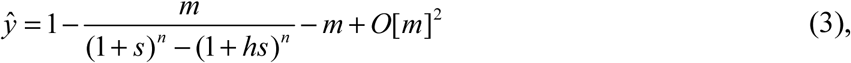

where *n* is the number of adaptive loci in inversion, which is opposite of the effect of *m* on the probability of establishment of a new inversion (see also Kirkpatrick and Barton 2006).

**Fig. 2.**
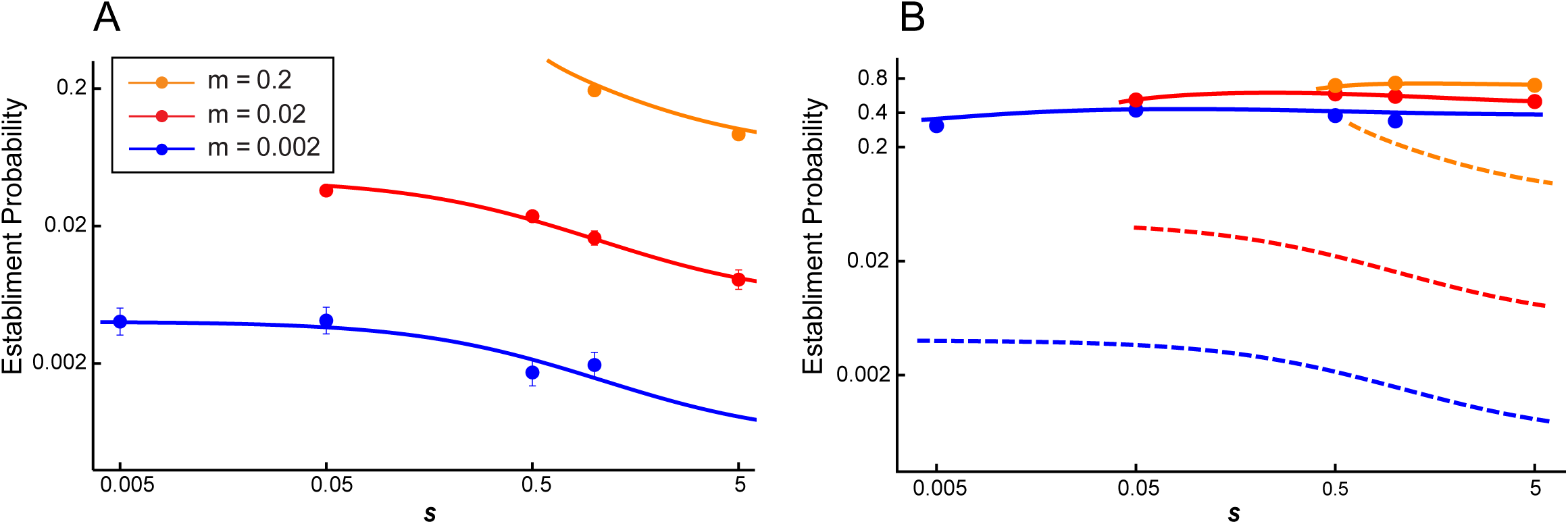
Comparison between establishment probability of new mutations versus that from standing inversion variation. (A) Establishment probability of a single new mutation of inversion in the marginal population at migration-selection balance for a population size (2*N)* of 10,000 under different orders of gene flow rate (*m*) and allele effect size (*s*). (B) Establishment probability of standing inversion variations (solid lines) compared with that from a single inversion mutation (dashed lines). Lines are theoretical predictions while solid circles are simulation results with 95% confidence levels shown as error bars.

### Probability of establishment of adaptive standing inversion variation

Now let us consider an adaptive inversion, *AB\**, that is segregating in a population at frequency *y* (Fig. 1C). When gene flow starts, with regards to the rate of increase in the frequency of the inversion, Eq.1 still holds, except that the frequencies of genotypes are not in equilibrium. Therefore, the rate of change of the inversion frequency becomes

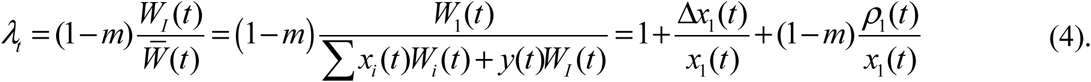

Somewhat surprisingly, following the influx of maladapted genotypes with the initiation of migration between the populations, our results show that the frequency of the inversion will actually decrease for a few generations (*λ_t_*<1). This is because gene flow will initially decrease the frequency of *x*_1_ (i.e., Δ*x*_1_ is negative), and with low frequencies of recombinants (*Ab* or *aB*) *ρ*_1_(*t*) is small (i.e., there is a small change in the frequency of *x*_1_ attributable to recombination with either locally coadapted or maladaptive alleles at time *t*). However, as the frequency of recombinants increases, the selective advantage of an inversion is realized when divergence occurs with gene flow. Thus, the probability that any single copy of segregating inversion surviving till generation *t*, *U_t_*, can be determined by integration of the changes in *λ* of each generation using a time heterogeneous branching process (Ohta and Kojima 1968), 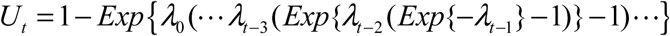. The probability of establishment of a segregating inversion for a given frequency *y* is just the probability that at least one copy of the inversion survives stochastic loss, Π*_y_* = 1 − (1 − *U*_∞_)^2*Ny*^. Since the segregating inversion can be viewed as neutral mutations prior to the onset of migration (Fig. 1C), the probability of observing *k* copies of inversions in a population 2*N* at the time when gene flow starts can be approximated as 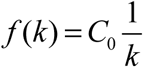, where 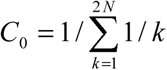 (Ewens 2004), assuming no back mutation.

Thus, the probability of establishment from standing inversion variation becomes

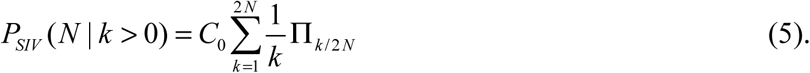

Considering a range of allele effect sizes, we show that the highest probability of establishment of segregating inversion variation occurs when selection is an order higher than the migration rate (Fig. 2B). Compared to the establishment probability of a new inversion, the probability of establishment of a segregating inversion (i.e., *k*>0) depends much more weakly on the migration rate *m* than in the case of new mutations (i.e., the establishment probability is logarithmically, not linearly, related to *m*; Fig. 2B).

### Comparison of the probability of adaptation from two sources of inversion variation

To determine the conditions under which adaptation from new mutations versus standing inversion variation is more probable, we have to consider not only the probability of establishment of the inversion (as discussed in the previous section), but also the availability of inversions. For new inversions, the relevant factors determining the availability of inversions is mutational input, whereas for standing genetic variation, the key parameter is the frequency distribution of segregating inversion variation upon the start of gene flow.

The input of new inversion mutations can be approximated as *θ* = 2*N_e_μ_I_x*_1_, where *x*_1_ is the frequency of coadapted genotype *AB* in the population and *μ_I_* is the mutation rate of inversions per gamete per generation that encompass the region of the chromosome where adaptive loci are located. Therefore, the probability of adaptation from a new inversion mutation within *T* generations is

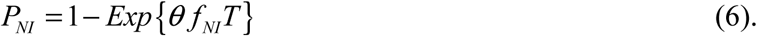

Similar to the establishment probability (*f_NI_*) for a new inversion mutation conditioned on the availability of a new mutation, *P_NI_* also increases along with *m* (Fig. 3A). The ratio of *s* relative to *m*, rather than exact values of s, is key to determining *P_NI_* given same *m*. While *f_NI_* is greatest at low values of *s* (Fig. 2A), when the waiting time for a new inversion mutation is taken into account, the probability of adaptation, *P_NI_*, is actually improbable at lower range of *s/m* (Fig. 3A). This is because when alleles are under weak selection, the frequency of the adaptive genotype *AB* is so low that it is unlikely for a new inversion mutation to capture it. Instead, the highest probability of adaptation is maximized at a moderate ratio of *s/m* because of the tradeoff between the rate of establishment and inversion availability (Fig. 3A).

**Fig 3.**
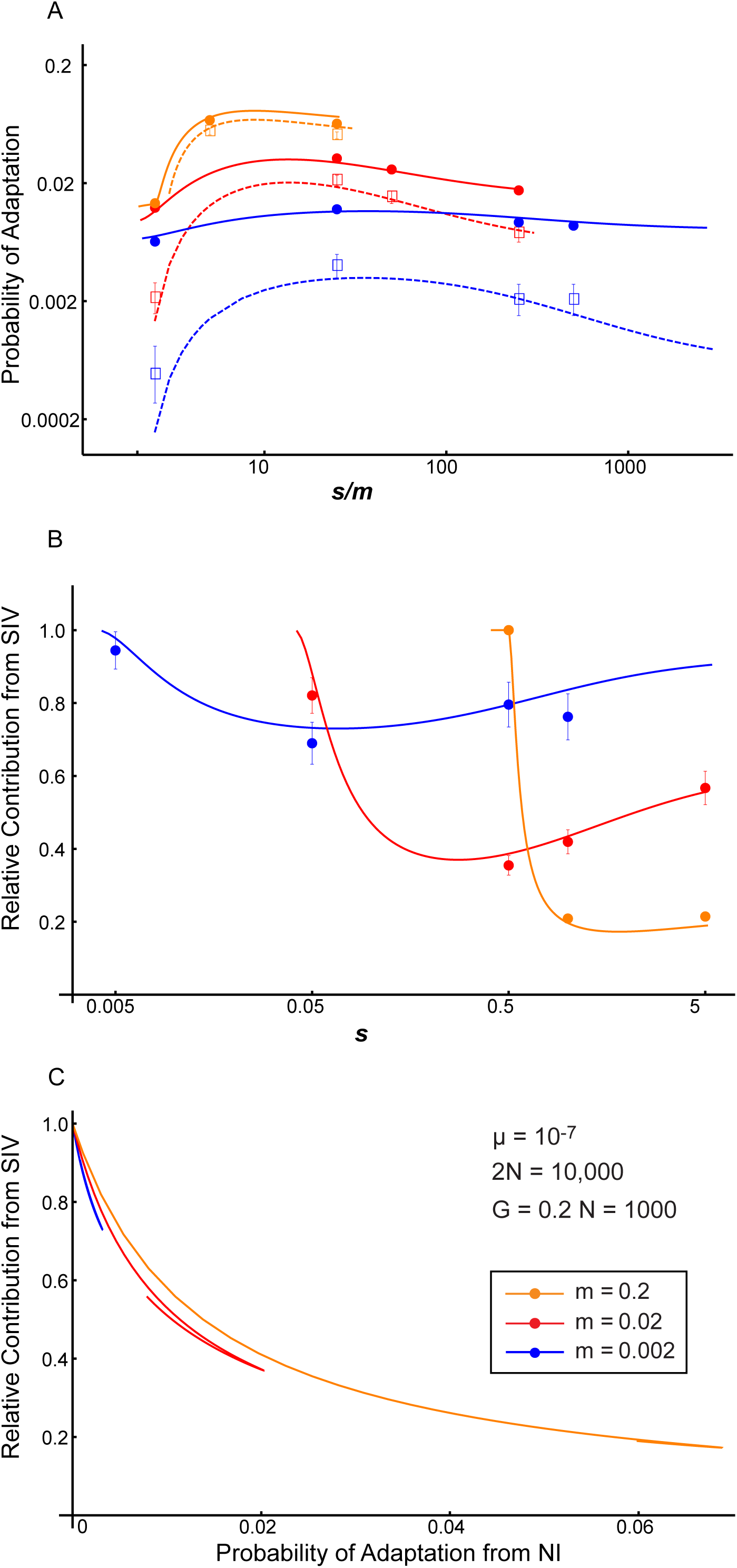
Probability of adaptation and relative contribution from standing variation. (A) Comparison between adaptation from new inversions (*P_NI_*; dashed lines) and adaptation from both sources (*P_ADP_*; solid lines) given new input of mutations persisting for *G* = 0.2*N_e_* generations after the initiation of maladaptive alleles (see Fig. 1) for a population size (2*N)* of 10,000 under different orders of *m* and *s*. Mutation rate, *μ*=10^-7^. 95% confidence levels of each simulation are showed on error bars of the squares (*P_ADP_*;) or circles (*P_NI_*). Note that the probability of adaptation is plotted against *s/m*. (B, C) Relative contribution of standing inversion variation to rapid divergence (i.e., within 0.2 *N* generations). (B) is plotted against *s*, (C) is plotted against *P_ADP_*.

Below we derive the probability of adaptation from standing inversion variation by integrating over the availability of the inversion and its establishment probability,

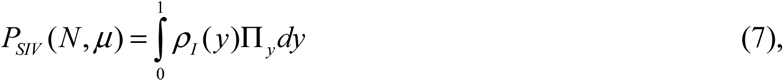

where the frequency spectrum of segregating inversions in the population at mutation-drift balance before the onset of migration can be derived (see Ewens 2004; Hermisson and Pennings 2005) as 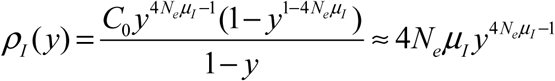, where *C_0_* is a constant of integration.

The shape of *P_SIV_*(*N*, *μ*) and *P_SIV_*(*N* | *k*> 0)for different values of *m* and *s* are similar (Fig. 2B, 3A), but *P_SIV_*(*N*, *μ*) scales with the availability of inversions, *N_eμI_*. This means the chance of observing standing inversion variation at a given time in the population is proportional to the mutation rate. In contrast with the establishment probability, which is always higher for standing inversion variation compared to new inversions, when the availability of the inversion is also considered, the probability of adaptation via standing inversion variation is not necessarily going to be higher than the probability of adaptation by new inversions.

### Contribution of standing inversion variation to adaptive divergence

For rapid adaptation via inversions under a divergence with gene flow model, how important is standing inversion variation relative to new inversion mutations as the likely source? This question can be evaluated by calculating the relative contribution of standing inversion variation to adaptive divergence, as derived by a combination of Eq. 6 and 7,

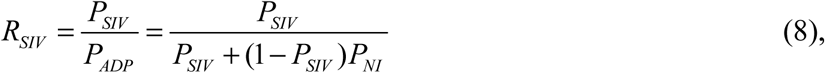

following ref. (Hermisson and Pennings 2005). As *P_NI_* increases with time, if time allowed for inversions to occur is long enough, *P_NI_* will eventually surpass *P_SIV_* regardless of the scenario. However, in this paper we are interested in rapid rescuing effect from inversions after secondary contact, we only simulate the situation when the time allowed for adaptation is short (i.e., 0.2 *N_e_* generations) so that source of inversion variation is highly relevant to the probability of adaptation.

There are several parameter regions where the relative contribution from standing inversion variation is particularly important (Fig. 3B), specifically, when *s* is at the same order of *m* so that *s* is too small to withstand the gene flow (see *m* = 0.2, *s* = 0.5 on Fig. 3A), or when allele effect sizes are large enough compared to migration (s>>m). These regions, however, all correspond to situations where adaptation via inversions are less to probable to occur. Plotting *R_SIV_* against *P_NI_* (Fig. 3C), we can see that as adaptation via new inversions becomes more probable (higher *P_NI_*), contribution from standing inversion variation quickly drops.

## Discussion

Both standing genetic variation and chromosomal inversions have become central foci as mechanisms to facilitate rapid adaptation (Barrett and Schluter 2008). By developing an analytical model that makes explicit the factors governing the dynamics of rapid adaptation based on inversion variation, we show that when adaptive divergence via inversions with gene flow is more likely, new mutational input (i.e., new inversion variation) becomes a more probable genetic source than standing inversion variation. By considering a broad range of selective values of alleles (instead of assuming weak selection (Kirkpatrick and Barton 2006)), we also use our model and simulation to predict (i) the characteristics of inversions contributing to adaptation (e.g., the selective benefit of alleles and its relationship with migration and number of loci involved) and (ii) conditions when the relative importance of segregating inversion variation as a source of rapid adaptation might be increased.

### Implications of results for the genetics of adaptation

Inversions are more likely to facilitate local adaption under higher gene flow rates (Fig 2; see also Kirkpatrick and Barton 2006). Consequently, we can identify how the genes contained within the inversion are likely to be involved in adaptation because of their impact on the effective gene flow rate. The ratio between allele effect size and gene flow, rather than the absolute value of the effect size, determines the likelihood of this scenario. The highest probability of adaptation is maximized at a moderate ratio because of the tradeoff between the rate of establishment and inversion availability (Fig. 3A). This finding predicts the genomic profile of adaptation, that is, whether divergence is achieved through multifarious selection on many genes or through linked regions within genetic islands (Nosil et al. 2009b, Nosil, 2012 #537) with inversions involved. The allele effect sizes of selected loci that can benefit the most from being captured in inversions will differ given different levels of gene flow. Under adaptation with strong gene flow (i.e., when population migration rate, *2Nm*, is much larger than 1 (Wright 1931)), multifarious selection on many small effect genes cannot resist gene flow effectively, leading to clustering of few genes with larger effects in freely recombining region (Yeaman and Whitlock 2011) (see Fig. 2 for increasing minimum *s* to withstand gene flow at higher *m*). In this case, it is beneficial for selected loci with even big allele effect sizes to be captured in inversions. On the other hand, under adaptation with weak gene flow, multifarious selection can be seen more often in freely recombining region. In this case, much smaller effect alleles would be found more often within inversions.

Although it is widely recognized that either increasing the recombination rate between loci, *r*, or number of adaptive loci, *n*, will affect the probability of adaptation (Table S1 and Eq. 2 and 3 in Kirkpatrick and Barton 2006), their interaction generates different expectations for the genetics of adaptation, because *r* and *n* are usually negatively correlated for a given length of inversion. In other words, capturing more adaptive loci within the same length of an inversion will continuously increase its selective advantage until the point when all of the fitness-related loci are tightly linked (Fig. S1). Therefore, the length of an inversion influences its fitness because longer inversions can capture more adaptive loci without tight linkage while shorter inversions are less likely to be advantageous. This result helps to explain the observed size distribution of inversions in natural populations. For example, in *Anopheles gambiae*, while rare chromosomal inversions were found to vary randomly in length (Pombi et al. 2008), common inversions which are more widely spread in the populations tend to be long.

### Contribution of standing inversion variation versus new inversions to adaptation

High initial frequency and the immediacy of standing genetic variation are frequently cited as reasons why it is a more probable source for rapid adaptation than new mutations (Barrett and Schluter 2008). As with adaptation via new point mutation versus standing genetic variation (see Innan and Kim 2004; Orr and Unckless 2008; Przeworski et al. 2005), standing inversion variation also has a significantly higher establishment probability (Fig. 2) by virtue of a higher segregating frequency in a population (i.e., they are not as sensitive to stochastic loss by genetic drift compared with new inversions). However, consideration of the establishment probability alone (e.g., Feder et al. 2011; Kirkpatrick and Barton 2006) is not sufficient for understanding the contribution of standing inversion variation relative to new inversions. As modeled here, the availability of inversions is critical for evaluating whether adaptation is actually probable (Fig. 3A). For point mutations, their availability is only determined by the effective population size and mutation rate, whereas it is more complicated for inversions. Therefore, the impact of the immediacy of segregating inversions on the relative contribution of standing inversion variation and new inversions to adaptive divergence varies under different scenarios (i.e., combinations of *m* and *s*).

Our results show that higher gene flow rates result in greater contributions of adaptation from standing inversion variation if probability of adaptation (*P_NI_*) is controlled for (Fig. 3C). This can be understood by considering that an inversion does not confer a fitness difference directly, in contrast with point mutations that facilitate rapid adaptation under sudden environmental change (Orr and Betancourt 2001; Przeworski et al. 2005), bottlenecks (Hermisson and Pennings 2005; Orr and Unckless 2008) or domestication events(Innan and Kim 2004). For inversions, the sudden influx of maladaptive alleles from migrants gives inversions an advantage over non-inverted chromosomes. Gene flow is not only the driving force of adaptation via inversions (i.e., the level of gene flow determines the selective advantage of inversions), but it also impacts the availability of inversions. Higher gene flow will lower the population size of favorable genotypes, making it less likely that inversion mutations will capture adaptive alleles. Under this scenario, using an available pool of standing inversion variation that already captured good genotypes becomes more important.

We also find the expected contribution of standing inversion variation to adaptive divergence decreases as the conditions become more favorable to adaptation via inversions (i.e., *P_NI_* increases; Fig. 3C). This is mainly because the probability for adaptation from standing inversion variation increases slower than that via new inversions when conditions become favorable (compare solid lines with dotted lines in Fig. 3A), such that standing inversion variation can only compensate for unfavorable situations (i.e., increase the probability of adaptation relative to new mutation when adaptive divergence is not likely), but not outcompete new inversions under favorable situations (see also Hermisson and Pennings 2005). The reasons are two-fold. First, the advantage of a higher initial frequency of standing variation levels off when the probability of establishment from a single copy increases with higher gene flow and moderate allele effect size, the very conditions when adaptive divergence via inversions is actually likely. Second, the selective advantage of segregating inversions gradually builds up as the frequency of favorable genotypes drops and recombinants are accumulated (*λ* changing from negative to positive in Eq. 4). In contrast, a new inversion in a population at migration-selection balance realizes its maximum selective advantage, giving it a higher survival rate compared to a pre-existing inversion.

### If adaptation occurs from standing inversion variation, what can we infer about the process of adaptation?

Although for the conditions explored here, standing inversion variation is less important overall when conditions are favorable for adaptation via inversions, this finding does not eliminate the potential importance of standing inversion variation. With an understanding of the relevant factors impacting the probability of adaptation from standing inversion variation, we can identify the evolutionary context where standing inversion variation is predicted to contribute to adaptation, as illustrated by the specific scenarios discussed below.

#### Time until establishment

Whether adaptation can occur rapidly may determine the likelihood of evolution change (Hermisson and Pennings 2005; Lynch 2010). Establishment times are consistently shorter for standing inversion variation as compared to new inversions (compare circles to squares in Fig. 4). This discrepancy will be even larger if the waiting time for a new mutation to occur is included. Consequently, given an equal probability of adaptation for new inversions and standing inversions (*P_NI_* = *P_SIV_*), a faster establishment rate (shorter establishment time) of standing inversion variation alone will be highly likely to lead to rapid local adaptations. This was empirically supported by the case of *Drosophila subobscura*, which has an establishment time for standing inversion variation as short as 25 years to reach a similar latitudinal cline of adaptive inversion polymorphism seen in the old world after introduction into New World in the early 1980s (Balanya et al. 2003). These inversions are shown to harbor favorable combinations of alleles (Rego et al. 2010; Santos 2009).

**Fig 4.**
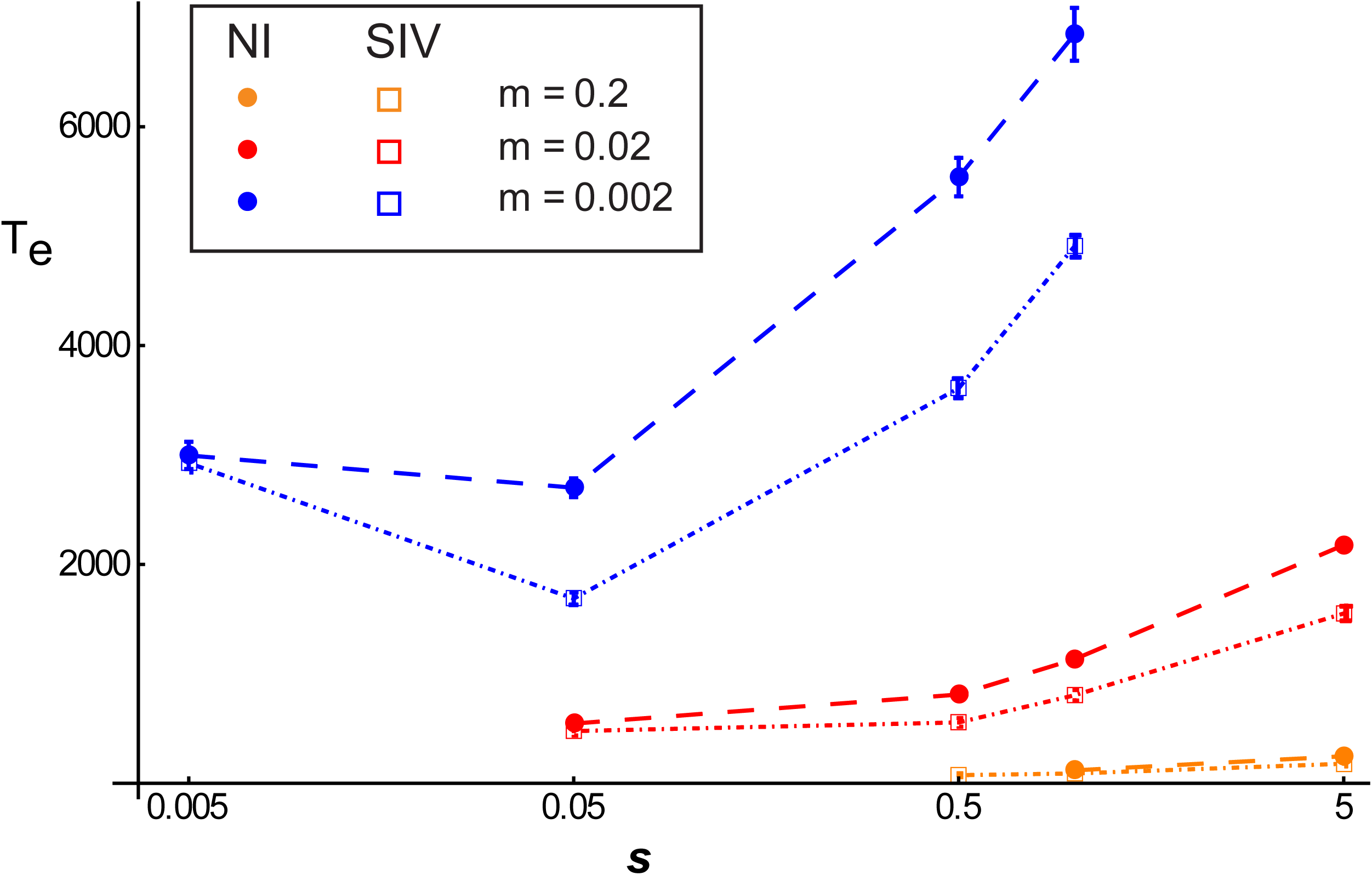
Average establishment time of inversions for new mutations versus standing variation calculated from runs with established inversions. Circles are waiting time for new inversions while squares are that for standing variations. Standard errors are shown as bars.

#### When and how standing inversion variation is introduced

While our findings hold when inversion variation evolves *de novo* within a focal population (where mutation rate sets the waiting time for new inversions as well as the chance of having segregating inversions), adaptive inversions introduced from populations located in similar environments could alleviate the recombination load that would accumulate in a population experiencing an influx of maladapted alleles from populations in dissimilar environments. Likewise, introgression from closely-related species is also a possible source of standing inversion variation. For example, the origin of the 2La and 2Rb inversions associated with dry environments in *Anopheles gambiae* (Coluzzi et al. 2002; White et al. 2007) trace back to an introgression event with *Anopheles arabiensis* (Besansky et al. 2003; Besansky et al. 1994). In another example of gene flow of adaptive inversions between populations, northerly distributed *Rhagoletis pomonella* gained inversion polymorphisms from Mexican populations that were strongly associated with the length of overwintering pupal diapauses, which facilitated a host shift (Feder et al. 2003b).

#### Genetic background

In our theoretical model, we assume that segregating inversions and new inversion mutations have the same fitness – that is, we do not consider the genetic background upon which the inversion occurs. Each new inversion mutation or segregating inversion can have a range of fitness values based on the genes they captured (Nei et al. 1967). Standing inversion variation may, in general, represent a more likely source of adaptive divergence because it will have already been exposed to purifying selection. The pre-filtering process would greatly decrease the frequency of inversions that capture genes with large fitness costs or have a direct fitness cost through meiotic problems. In other words, inversions with a lower fitness cost will segregate at a higher frequency, in contrast to new inversion variants, which have yet to pass through the selection gauntlet.

#### Demographic histories

Adaptive divergence in empirical populations may of course occur under conditions other than the constant population sizes modeled here. For example, population sizes may fluctuate, especially in response to shifts in climatic or ecological factors. Such changes in population sizes are relevant because they will not only influence the amount of mutational input but also the relative contribution of new inversions versus standing inversion variation to adaptive divergence (Hermisson and Pennings 2005; Kimura and Crow. 1970; Orr and Unckless 2008; Otto and Whitlock 1997). To compare the relative contributions of new inversions versus standing inversion variation with fluctuating population sizes, we considered a scenario involving cyclic dynamics (such as those observed in mosquito populations) where population sizes differ depending on climatic conditions (e.g., wet/dry or warm/cold). We did forward-time simulations using parameters selected from empirical studies of *Anopheles gambiae* populations (Manoukis et al. 2008), which are characterized by multiple inversion polymorphisms. We show that when the probability of adaptation becomes larger under different combinations of *m*, *s*, *r*, and *n*, the relative importance of standing inversion variation also decreases with fluctuating population sizes (Fig. 5), which is similar to what has been demonstrated under theoretical predictions (Fig. 3C). However, population fluctuations affect the steepness of the negative relationship between probability of adaptation and contribution from standing variation. When a population is cyclic, contribution from standing inversion variation is higher (Fig. 5). This is contingent on the assumption that the onset of gene flow occurs at the time when population size is increasing, in order to mimic the situation where populations begin to multiply and migrate when the wet season begins. Therefore, if inversions are pre-existing, they have much less chance to be lost by drift. The situation will be reversed when gene flow occurs in a shrinking population. However, the first situation is more probable under the context of secondary contact. In either case, the proportional contribution from standing inversion variation should be boosted or decreased by a factor of *N/N*_e_ according to (Otto and Whitlock 1997).

**Fig 5.**
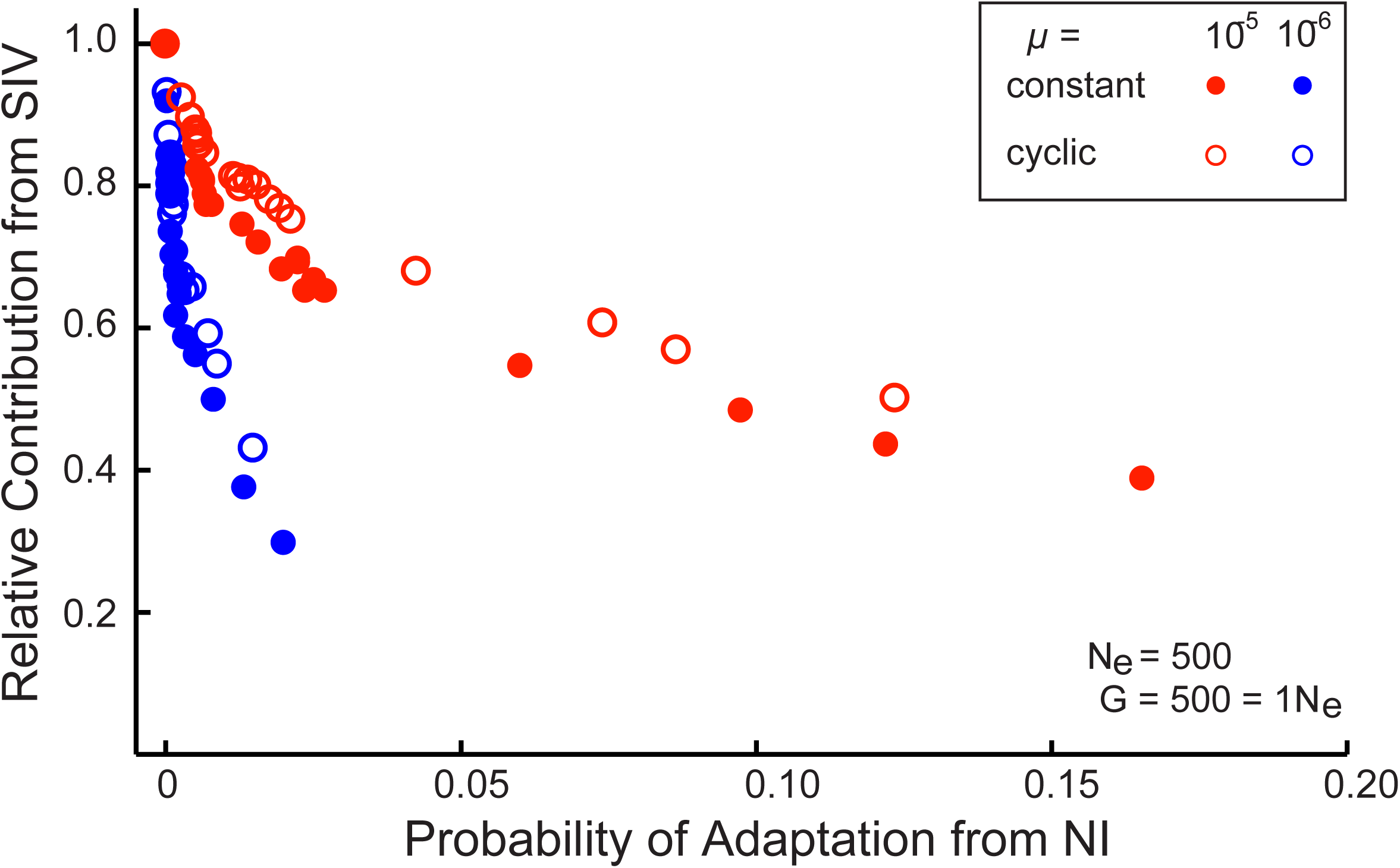
The relationship between probability of adaptation from new inversion and relative contribution from standing inversion variation (*N_e_* = 500, *G=1N_e_*) for different parameter settings (*m*, *s*, *r*, *n)* under two demographic scenarios: a constant population (*N* = *N_e_* = 500) and cyclic population (*N*(*t*) = 2525 + 2475sin(2*π*(*t* + 6.5) /10)). New input of mutations lasts for 500 generations (*G* = 1*N*), with two levels of mutation rate, 10^-6^ and 10^-5^. 5,000 realizations in each scenario were run to observe the impact of different combination of parameters on the proportion of contribution from standing variation to the success of establishment of inversions. Blue colored and red colored circles denote different level of mutation rate, 10^-6^ and 10^-5^, respectively. Open circles are simulation results from cyclic population while filled circles are from constant population realizations.

## Acknowledgements

We appreciate useful discussions with Tim Connallon, Trevor Bedford, Huateng Huang, and Mark Kirkpatrick.

## References

Balanya, J., L. Serra, G. W. Gilchrist, R. B. Huey, M. Pascual, F. Mestres, and E. Sole. 2003. Evolutionary pace of chromosomal polymorphism in colonizing populations of *Drosophila subobscura*: An evolutionary time series. Evolution 57:1837–1845.

Barrett, R. D. H., and D. Schluter. 2008. Adaptation from standing genetic variation. Trends in Ecology & Evolution 23:38–44.

Besansky, N. J., J. Krzywinski, T. Lehmann, F. Simard, M. Kern, O. Mukabayire, D. Fontenille et al. 2003. Semipermeable species boundaries between *Anopheles gambiae* and *Anopheles arabiensis*: Evidence from multilocus DNA sequence variation. Proceedings of the National Academy of Sciences of the United States of America 100:10818–10823.

Besansky, N. J., J. R. Powell, A. Caccone, D. M. Hamm, J. A. Scott, and F. H. Collins. 1994. Molecular phylogeny of the *Anopheles gambiae* complex suggests genetic introgression between principal malaria vectors. Proceedings of the National Academy of Sciences of the United States of America 91:6885–6888.

Bradshaw, H. D., S. M. Wilbert, K. G. Otto, and D. W. Schemske. 1995. Genetic-mapping of floral traits associated with reproductive isolation in monkeyflowers (*Mimulus*). Nature 376:762–765.

Colosimo, P. F., K. E. Hosemann, S. Balabhadra, G. Villarreal, M. Dickson, J. Grimwood, J. Schmutz et al. 2005. Widespread parallel evolution in sticklebacks by repeated fixation of ectodysplasin alleles. Science 307:1928–1933.

Coluzzi, M., A. Sabatini, A. della Torre, M. A. Di Deco, and V. Petrarca. 2002. A polytene chromosome analysis of the *Anopheles gambiae* species complex. Science 298:1415–1418.

Ewens, W. J. 2004, Mathematical population genetics. I. Theoretical Introduction: Interdisciplinary Applied Mathematics. New York, Springer.

Feder, J. L., S. H. Berlocher, J. B. Roethele, H. Dambroski, J. J. Smith, W. L. Perry, V. Gavrilovic et al. 2003a. Allopatric genetic origins for sympatric host-plant shifts and race formation in *Rhagoletis*. Proceedings of the National Academy of Sciences of the United States of America 100:10314–10319.

Feder, J. L., R. Gejji, T. H. Q. Powell, and P. Nosil. 2011. Adaptive chromosomal divergence driven by mixed geographic mode of evolution. Evolution 65:2157–2170.

Feder, J. L., F. B. Roethele, K. Filchak, J. Niedbalski, and J. Romero-Severson. 2003b. Evidence for inversion polymorphism related to sympatric host race formation in the apple maggot fly, *Rhagoletis pomonella*. Genetics 163:939–953.

Fisher, R. A. 1930, The genetical theory of natural selection. Oxford, U.K., Oxford Univ. Press.

Haldane, J. B. S. 1927. A mathematical theory of natural and artificial selection. Part V. Selection and mutation. Proceedings of the Cambridge Philosophical Society 23:838–844.

Hermisson, J., and P. S. Pennings. 2005. Soft sweeps: Molecular population genetics of adaptation from standing genetic variation. Genetics 169:2335–2352.

Innan, H., and Y. Kim. 2004. Pattern of polymorphism after strong artificial selection in a domestication event. Proceedings of the National Academy of Sciences of the United States of America 101:10667–10672.

Joron, M., L. Frezal, R. T. Jones, N. L. Chamberlain, S. F. Lee, C. R. Haag, A. Whibley et al. 2011. Chromosomal rearrangements maintain a polymorphic supergene controlling butterfly mimicry. Nature 477:203–206.

Karasov, T., P. W. Messer, and D. A. Petrov. 2010. Evidence that adaptation in *Drosophila* is not limited by mutation at single sites. PLoS Genet 6:e1000924.

Kimura, M. 1957. Some Problems of Stochastic Processes in Genetics. The Annals of Mathematical Statistics 28:882–901.

Kimura, M. 1983, The neutral theory of molecular evolution. Cambridge, U.K., Cambridge Univ. Press.

Kimura, M., and J. F. Crow. 1970, An introduction to population genetics theory. New York, Harper & Row.

Kirkpatrick, M. 2011. How and why chromosome inversions evolve. Plos Biology 8:5.

Kirkpatrick, M., and N. Barton. 2006. Chromosome inversions, local adaptation and speciation. Genetics 173:419–434.

Lewontin, R. C., and K. Kojima. 1960. The evolutionary dynamics of complex polymorphisms. Evolution 14:458–472.

Li, W. H., and M. Nei. 1974. Stable linkage disequilibrium without epistasis in subdivided population. Theoretical population biology 6:173–183.

Lowry, D. B., and J. H. Willis. 2010. A widespread chromosomal inversion polymorphism contributes to a major life-history transition, local adaptation, and reproductive isolation. Plos Biology 8:14.

Lynch, M. 2010. Scaling expectations for the time to establishment of complex adaptations. Proceedings of the National Academy of Sciences of the United States of America 107:16577–16582.

Manoukis, N. C., J. R. Powell, M. B. Toure, A. Sacko, F. E. Edillo, M. B. Coulibaly, S. F. Traore et al. 2008. A test of the chromosomal theory of ecotypic speciation in *Anopheles gambiae*. Proceedings of the National Academy of Sciences of the United States of America 105:2940–2945.

Navarro, A., and N. H. Barton. 2003. Accumulating postzygotic isolation genes in parapatry: A new twist on chromosomal speciation. Evolution 57:447–459.

Nei, M., K.-I. Kojima, and H. E. Schaffer. 1967. Frequency changes of new inversions in populations under mutation-selection equilibria. Genetics 57:741–750.

Noor, M. A. F., K. L. Grams, L. A. Bertucci, and J. Reiland. 2001. Chromosomal inversions and the reproductive isolation of species. Proceedings of the National Academy of Sciences of the United States of America 98:12084–12088.

Nosil, P., D. J. Funk, and D. Ortiz-Barrientos. 2009a. Divergent selection and heterogeneous genomic divergence. Molecular Ecology 18:375–402.

Nosil, P., L. J. Harmon, and O. Seehausen. 2009b. Ecological explanations for (incomplete) speciation. Trends in Ecology & Evolution 24:145–156.

Ohta, T., and K. I. Kojima. 1968. Survival probabilities of new inversions in large populations. Biometrics 24:501–516.

Orr, H. A. 1998. The population genetics of adaptation: The distribution of factors fixed during adaptive evolution. Evolution 52:935–949.

Orr, H. A., and A. J. Betancourt. 2001. Haldane’s sieve and adaptation from the standing genetic variation. Genetics 157:875–884.

Orr, H. A., and R. L. Unckless. 2008. Population extinction and the genetics of adaptation. American Naturalist 172:160–169.

Otto, S. P., and M. C. Whitlock. 1997. The probability of fixation in populations of changing size. Genetics 146:723–733.

Papadopulos, A. S. T., W. J. Baker, D. Crayn, R. K. Butlin, R. G. Kynast, I. Hutton, and V. Savolainen. 2011. Speciation with gene flow on Lord Howe Island. Proceedings of the National Academy of Sciences of the United States of America 108:13188–13193.

Pombi, M., B. Caputo, C. Costantini, M. A. Di Deco, M. Coluzzi, A. della Torre, F. Simard et al. 2008. Chromosomal plasticity and evolutionary potential in the malaria vector *Anopheles gambiae sensu stricto*: insights from three decades of rare paracentric inversions. BMC evolutionary biology 8:309.

Przeworski, M., G. Coop, and J. D. Wall. 2005. The signature of positive selection on standing genetic variation. Evolution 59:2312–2323.

Rego, C., J. Balanya, I. Fragata, M. Matos, E. L. Rezende, and M. Santos. 2010. Clinal patterns of chromosomal inversion polymorphisms in *Drosophila subobscura* are partly associated with thermal preferences and heat stress resistance. Evolution 64:385–397.

Rieseberg, L. H. 2001. Chromosomal rearrangements and speciation. Trends in Ecology & Evolution 16:351–358.

Santos, M. 2009. Recombination Load in a Chromosomal Inversion Polymorphism of *Drosophila subobscura*. Genetics 181:803–809.

Scoville, A. G., and M. E. Pfrender. 2010. Phenotypic plasticity facilitates recurrent rapid adaptation to introduced predators. Proceedings of the National Academy of Sciences of the United States of America 107:4260–4263.

White, B. J., M. W. Hahn, M. Pombi, B. J. Cassone, N. F. Lobo, F. Simard, and N. J. Besansky. 2007. Localization of candidate regions maintaining a common polymorphic inversion (2La) in *Anopheles gambiae*. PLoS Genet 3:e217.

Wright, S. 1931. Evolution in Mendelian populations. Genetics 16:0097–0159.

Yeaman, S., and M. C. Whitlock. 2011. The genetic architecture of adaptation under migration-selection balance. Evolution 65:1897–1911.

